# Effects of transcranial direct current stimulation on brain networks related to creative thinking

**DOI:** 10.1101/2020.03.08.981506

**Authors:** Koji Koizumi, Kazutaka Ueda, Ziyang Li, Masayuki Nakao

## Abstract

Human creative thinking is unique and capable of generating novel and valuable ideas. Recent research has clarified the contribution of different brain networks (default mode network, DN; executive control network; salience network) to creative thinking. However, the effects of brain stimulation on brain networks during creative thinking and on creative performance have not been clarified. The present study was designed to examine the changes in functional connectivity (FC) and effective connectivity (EC) of the large-scale brain network, and the ensuing changes in creative performance, induced by transcranial direct current stimulation (tDCS). Fourteen healthy male students underwent two tDCS sessions, one with actual stimulation and one with sham stimulation, on two separate days. Participants underwent tDCS (anode over the left dorsolateral prefrontal cortex, DLPFC; cathode over the right inferior parietal lobule, IPL) for 20 minutes. Before and after the tDCS session, electroencephalography signals were acquired from 32 electrodes over the whole head during the creative thinking task. On FC analysis, the delta band FC between the posterior cingulate cortex and IPL significantly increased only after real stimulation. We also found that the change of flexibility score was significantly correlated with the change in (i) delta band FC between mPFC and left lateral temporal cortex (LTC) and (ii) alpha band FC between IPL and right LTC. On EC analysis, decreased flow within the DN (from left LTC to right IPL) was observed. Our results reveal that tDCS could affect brain networks, particularly the DN, during creative thinking and modulate key FC in the generation of flexible creative ideas.

## 1 Introduction

Science, technology, education, and culture, the things that permeate and enrich every part of our lives, are the products of creativity, which is humankind’s ultimate resource (Toynbee, 1964). However, no comprehensive definition of creativity has been achieved yet, because creativity is a multifaceted construct formed by different strands pertaining to the person who creates, the mental processes of creating ideas, and the influence of the environment on the person and the product as an outcome (Rhodes, 1961). Here, we borrowed the most accepted definition of creativity, the ability to generate knowledge or work that is novel and useful (Barron, 1955; Sternberg and Lubart, 1996; Runco and Jaeger, 2012; Diedrich et al., 2015). To date, the main research interest in the field of neuroscience of creativity are the mental processes of creative thinking involving both the generation of novel ideas and the evaluation or selection of useful ones. Generation refers to coming up with various novel ideas outside the box and/or providing various solutions in response to an open-ended problem or divergent thinking task. Evaluation, on the other hand, refers to testing solutions by logical thinking and reasoning from known information, and then selecting the correct useful option in response to a closed-ended problem or convergent thinking task. The alternative uses task (AUT) has been often used as a divergent thinking task to generate as many uncommon use ideas as possible for everyday objects. Such divergent thinking task could measure personal-psychological creativity, or the ability of generating ideas novel to the person who create them, irrespective of whether other people have had those ideas before (Boden, 2004; Gilhooly et al., 2007). Furthermore, a validation study suggested that divergent thinking ability is better correlated with real-world creative achievement such as inventions, or published articles than convergent thinking ability, or intelligence quotients (Plucker, 1999). Therefore, the present study focused on idea generating processes during divergent thinking as a key component of creative thinking.

Previous studies have revealed that creativity is a complex construct that requires the involvement of various brain functions such as memory, future simulation, semantic processing, and attention. Besides, recent neuroimaging studies of creativity have suggested the involvement of multiple brain networks, including the following: the default mode network (DN), the executive control network (ECN), and the salience network (SN) (Beaty et al., 2016, 2017, 2019; Yu-chu Yeh et al., 2019; Girn et al., 2020). The relationship between divergent thinking and brain regions within the DN has been reported in several fMRI studies (Fink et al., 2010; Takeuchi et al., 2012; Gonen-Yaacovi et al., 2013; Beaty et al., 2014; Wei et al., 2014). The core DN subsystem (DN_CORE_) consists of the medial prefrontal cortex (mPFC), posterior cingulate cortex (PCC), and bilateral inferior parietal lobule (IPL) and has been associated with spontaneous generation of creative ideas, as it was found to be involved in internally oriented tasks such as mind wandering, episodic memory retrieval, autobiographical future thinking (Mason et al., 2007; Buckner et al., 2008; Spreng et al., 2009; Andrews-Hanna, 2012). Takeuchi et al. (2012) reported that the divergent thinking test score positively correlated with the strength of resting-state functional connectivity between the mPFC and PCC. The DN subsystem (DN_MTL_), centered around the medial temporal lobe (MTL) and including the hippocampal formation (HF), has also been reported to get activated together with the DN_CORE_ during divergent thinking (Madore et al., 2019). DN_MTL_ involves semantic/episodic memory and constructive mental simulations (Addis et al., 2007; Schacter et al., 2012; Moscovitch et al., 2016; Duff et al., 2020). A number of studies have suggested the involvement of the semantic network in generating ideas (Kenett et al., 2014; Hass, 2017; Kenett, 2018; Beaty et al., 2020). When generating creative ideas on divergent thinking, the process of combining remote associations in a novel way through the semantic network is promoted. The hippocampus plays the important role of taking existing mental concepts or associative information out of the original context and combining them to into a new context in service of divergent thinking (Luo and Niki, 2003; Duff et al., 2013; Backus et al., 2016). The middle temporal gyrus (MTG) within the lateral temporal cortex (LTC, the third DN subsystem [DN_SUB3_]) also involves semantic processing (Wei et al., 2012) and plays a key role in distant conceptual association in creative insights (Shen et al., 2017). Previous studies suggested that MTG facilitates the integration of information in the DN (Davey et al., 2016) and the MTL’s ability to detect novel features by novel semantic associations (Ren et al., 2020). On the contrary, the ECN, mainly consisting of the VLPFC and the dorsolateral PFC (DLPFC) plays a key role in deliberate cognitive control, such as working memory, relational integration, attentional shift, and task switching (Dreher and Berman, 2002; Curtis and D’Esposito, 2003; Wager et al., 2004; Blumenfeld et al., 2011). Additionally, previous studies have suggested that the ECN plays important roles in generating creative ideas during divergent thinking, such as in semantic processing (Fink et al., 2009), maintaining an internally generated thought (Burgess et al., 2003), flexibly switching between semantic categories (Kleibeuker et al., 2013), combining stored information (Dietrich, 2004), and novel idea organization (Wu et al., 2015). The salience network (Salience), mainly consisting of the anterior insula (AI) and the anterior cingulate cortex (ACC), has been proposed to detect both external and internal salient events (Seeley et al., 2007). The ACC relates to the monitoring of competing information about choices between multiple association options and may activate the ECN, particularly the DLPFC which tends to selectively increase the weights of attributes relevant to task performance (Perlovsky and Levine, 2012). Several previous studies have suggested that the ACC also relates to the suppression of unwanted self-generated thoughts or memories from the DN_MTL_ with ECN (Anderson et al., 2004; Mitchell et al., 2007) and involves the semantic processing of remote associations (Howard-Jones et al., 2005). Wu et al. (2015) suggested the involvement of the right ACC in suppressing irrelevant thoughts or memories and monitoring and forming distant semantic associations during divergent thinking. Furthermore, previous fMRI studies have also investigated the cooperation and dynamic interactions of brain networks during divergent thinking using AUT (Beaty et al., 2015; Heinonen et al., 2016). Increased coupling between DN and SN was observed at the beginning, followed by increased coupling between the DN and ECN. A recent fMRI study focused on product-based creative thinking in which the DN and SN attenuated with time, whereas the activity of the ventrolateral prefrontal cortex (VLPFC) within ECN increased in later stages (Yu-chu Yeh et al., 2019).

As outlined above, most previous neuroimaging studies focused on correlational methods using electroencephalography (EEG) or fMRI and clarified the contribution of large-scale brain networks to creative thinking. Besides research on the neural correlates of creative thinking, several previous studies investigated the possibility of modulating brain networks related to creativity or enhancing it, using brain stimulation like transcranial direct current stimulation (tDCS) (reviewed in Lucchiari et al., 2018). Since current flows from the anode to the cathode, the tDCS acts directly on the pyramidal cells by flowing perpendicularly to the cortex just below the electrode. The mechanism of action of tDCS has been speculated to be based on the effect of DC current on the membrane potential. In other words, cortical excitability increases under the anode, and hyperpolarization under the cathode is thought to decrease excitability of the cell membrane (Schlaug and Renga, 2008; Antal and Herrmann, 2016). Recent previous research showed that the tDCS has an impact not only on target brain areas, but also on other areas and networks during both tasks and rest (Lang et al., 2005; Keeser et al., 2011; Polanía et al., 2011; Meinzer et al., 2012; Axelrod et al., 2015; Kajimura et al., 2016). Since creativity is a complex construct, the focus of each tDCS study’s interests and stimulation parameters (e.g., target creativity process, functions, brain region, etc.) has been varied. Previous studies focused on the inhibition mechanism trying to maintain thinking inside-the-box and reported the positive effects on divergent thinking by applying cathodal stimulation on the left inferior frontal gyrus (IFG) (Chrysikou et al., 2013; Hertenstein et al., 2019; Ivancovsky et al., 2019). In the research by Hertenstein et al. (2019), resting-state EEG activity was measured before and after applying tDCS on the IFG for further elucidation of the neural basis of creativity. These authors reported the increase of beta power of the right frontal area by anodal stimulation and its association with better performance. Colombo et al. (2015) reported that anodal stimulation over the left DLPFC increased the performance in a divergent thinking task but only after divergent priming, a result supporting the role of the ECN in attentional shift. Zmigrod et al. (2015) also applied anodal/cathodal stimulation over the left/right DLPFC during AUT and produced numerically higher creativity scores but did not reach significance levels. While there are a number of creativity studies that applied tDCS focusing on the inhibitory mechanism of IFG and the role of ECN, there are few tDCS studies that have focused on DN function, the core network of divergent thinking. Axelrod et al. (2015) reported that applying anodal tDCS on the left DLPFC (l-DLPFC) promotes mind wandering or task-unrelated thoughts and argued that this stimulation may indirectly affect the DN. In addition, anodal/cathodal tDCS of the left LPFC/right IPL increased mind wandering and the reverse electrode positions decreased mind wandering (Kajimura and Nomura, 2015). Considering the study report that mind wandering facilitates creative incubation and improves divergent thinking performance (Baird et al., 2012), anodal/cathodal stimulation of the l-DLPFC/ r-IPL might be promising for improvement of divergent thinking. Kajimura et al. (2016) suggested that applying tDCS on the left LPFC/right IPL (r-IPL) may affect the balance of the whole DN system, changing the attention focus and thus promoting or inhibiting imaginative processes. Furthermore, dynamic causal modeling analysis on resting-state found that the right IPL, but not the left, causally affects the activity of other DN regions (Di and Biswal, 2014). Considering these studies and the role of ECN in mediating the interactions between brain networks, we hypothesized that the anodal/cathodal stimulation of the l-DLPFC/ r-IPL could modulate the system balance of the large-scale brain network, especially the DN, causally affecting divergent thinking.

In the creativity research field, for the challenging goal of improving creativity, previous studies investigated tDCS effects on creativity performance and inferred the effects on brain function. However, while recent neuroimaging research has revealed the key involvement of multiple brain networks in creativity, there are few studies investigating the effects of tDCS on the large complex networks that form the basis of creativity, and how it affects performance as a cause and effect of network changes. Only by neuroimaging methods such as fMRI and EEG, the associations of brain activity with cognitive functions and behavior can be revealed. However, it is difficult to determine causality, in other words how brain activity affects cognitive functions and behavior. On the other hand, only by tDCS, it is difficult to determine whether performance changes are caused by the excited/inhibited areas or by a compensatory network mechanism maybe triggered by such perturbations. Luft et al. (2014) suggested the importance of combining brain stimulation methods with neuroimaging techniques and employing state-of-the-art connectivity data analysis techniques to obtain a deeper understanding of the underlying spatiotemporal dynamics of connectivity patterns and creative performance. In the present study, before and after applying tDCS, we measured EEG activity over the whole head during divergent thinking using AUT and selected core regions of brain networks related to creativity as ROIs and investigated the change of connection strength between ROIs. We also examined the correlation between changes in connectivity and changes in creativity scores to see how network connection changes causally impact creativity performance.

## 2 Materials and methods

### 2.1 Participants

Sixteen healthy male subjects with (1) no history of neurological or psychiatric disease, (2) no history of intracranial metal implantation, and (3) normal or corrected-to-normal vision participated in the experiment. The two participants with the highest sleepiness scores according to the Stanford Sleepiness Scale, assessed during the experiment, were excluded from the analysis considering the possibility that high sleepiness may affect brain activity and task performance during divergent thinking, and make the interpretation of results difficult. Previous studies (Wimmer et al., 1992; Horne, 1998) have shown that sleep loss negatively impacts the divergent thinking performance. Furthermore, a recent study (Vartanian et al., 2014) showed that sleep deprivation, producing greater levels of sleepiness, was associated with greater activation in the left inferior frontal gyrus (IFG) during an alternative uses task. The remaining 14 participants (23 ± 1.9 years old) were all right-handed. The research was approved by the Research Ethics Committee of the Graduate School of Engineering of the University of Tokyo (approval number: KE18-28) and was conducted in accordance with the Declaration of Helsinki. Participants were informed of the study purpose, and they provided their consent in writing. Since we do not have consent of the participants to publish the data, we cannot share it. The study was also conducted in accordance with the report of the Committee on Brain Stimulation in Japan (Ugawa et al., 2011) and internationally accepted safety standards (Nitsche et al., 2003b; Poreisz et al., 2007). The experiment was conducted in a laboratory within a few minutes walking distance from the University of Tokyo Hospital, and in case of any discomfort, it was possible to immediately stop the experiment, but no adverse event was observed.

### 2.2 A priori power analysis

We calculated the sample size based on a power analysis using G*Power version 3.1.9.7 (Faul et al., 2007, 2009). We determined the input parameters (effect size *dz* = 0.93, α error probability *p* = 0.05, and power = 0.8) and conducted two-tailed Wilcoxon signed-rank test. The determined effect size is the median effect size for nominally statistically significant results reported in recent meta-research (Szucs and Ioannidis, 2017) and higher than the optimistic lower bound (0.75) reported in a recent review article (Poldrack et al., 2017). The α error probability and power are standard levels for most fields including neuroimaging studies. A recent neuroimaging study (Song et al., 2019) also employed almost the same input parameters for sample size estimation in a priori-power analysis.

Results showed that a sample size of 12 was adequate to attain reliable effects. Therefore, we enrolled 14 participants.

### 2.3 tDCS

tDCS was applied to the participants’ scalp through a saline-soaked pair of sponge electrodes (5×7 cm) connected to a DC-Stimulator 1×1 tES device (Soterix Medical Inc, New York, USA). Electrodes were located according to the extended 10-20 international system for EEG, based on previous reports and the research on cortical projection of EEG sensors (Koessler et al., 2009; Axelrod et al., 2015; Kajimura et al., 2016). The anode was placed over F3, corresponding to the left DLPFC, and the cathode over P4, corresponding to the r-IPL. Real direct current stimulation (Real-Stim) consisted of a constant 2 mA current (current density: 0.057 mA/cm^2^) applied for 20 minutes with a 30-second fade-in and fade-out. Previous studies have suggested that cortical excitability is stable for at least 1 hour after a 20-minute stimulation (Nitsche and Paulus, 2000, 2001; Nitsche et al., 2003c, 2003a). Participants perceived the direct current as an itching sensation at the anode contact point at the beginning of the stimulation. To produce sham stimulation (Sham-Stim), the DC-Stimulator has a built-in placebo mode. The electrodes were placed in the same arrangement, but the current stimulation was only delivered for the 30-second ramp up and the 30-second ramp down at the beginning and the end, to mimic the somatosensory artifacts of real stimulation (DaSilva et al., 2011). In the sham condition, the participants received no current for the rest of the 20-minute stimulation period, thus experiencing the itching sensation only at the beginning. This procedure makes it possible to keep participants blind to their stimulation condition. In this study, no subject recognized that one stimulus condition was a sham.

### 2.4 Assessment of individual creativity performances

The paper-and-pencil version of the alternative uses task (AUT) (Guilford et al., 1978) was used. The AUT is widely used in creativity research and is meant to measure the participants’ ability for divergent thinking (Runco and Acar, 2012). In the AUT, an everyday object was presented, and the participants were instructed to write down as many alternative uses as possible for the object, different from its common use. Before the task, the participants were shown the alternative uses of a newspaper, as an example, and practiced writing down the alternative uses of a chair for one minute. We selected 20 objects frequently used in previous studies (Rossmann and Fink, 2010; Glenn Dutcher, 2012; Lee and Therriault, 2013; Zmigrod et al., 2015; Yamaoka and Yukawa, 2016) and performed four types of AUT (A, B, C, and D) for the four conditions (Pre/Post Real-Stim and Pre/Post Sham-Stim) (Table 1). Each type of AUT consisted of five objects with a time limit of two minutes per object, counterbalanced among participants.

**Table 1.**
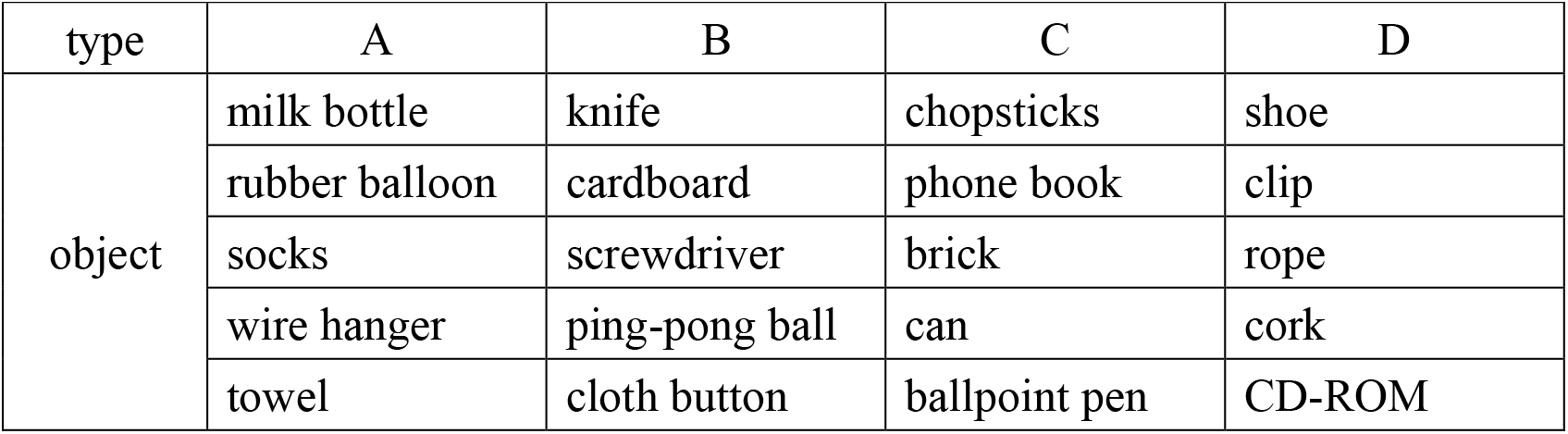
List of 20 objects used in the four types of the Alternative Uses Task (AUT)

| type | A | B | C | D |
| --- | --- | --- | --- | --- |
| object | milk bottle | knife | chopsticks | shoe |
|  | rubber balloon | cardboard | phone book | clip |
|  | socks | screwdriver | brick | rope |
|  | wire hanger | ping-pong ball | can | cork |
|  | towel | cloth button | ballpoint pen | CD-ROM |

Creativity was scored along three dimensions: fluency (the ability to produce a large number of ideas), flexibility (the ability to produce diverse categories of ideas), and originality (the ability to produce novel and unique ideas) (Guilford et al., 1978). Three university students (two males and one female) were hired as judges to rate the flexibility and originality. The creativity score of each condition was calculated as follows:

a. Fluency: average number of relevant answers for each everyday object.
b. Flexibility: The judges were instructed to evaluate the number of categories into which the answers for each everyday object could be classified. The average score of five objects evaluated by three judges was calculated as the task score. Intraclass coefficient (ICC) values for flexibility scores of all objects ranged between 0.53 and 0.97 for inter-rater reliability (Table 2). ICC values for each task type, which were the averaging values of the five objects, ranged from 0.69 to 0.89. These were good or excellent reliability levels based on the interpretation in the guidelines by Cicchetti (2001).
c. Originality: The judges rated the answers for each object regarding their novelty and uniqueness. Specifically, the judges were instructed to evaluate the originality of the answers on a five-point rating scale ranging from 1 (not original at all) to 5 (highly original) (De Dreu et al., 2008; Silvia et al., 2008; Fink et al., 2009) with the following criteria: “Whether the answer to a question hard to come up with or to imagine from the way we usually use things, was clever or not” based on the instructions for judging creativity by Silvia et al. (2008). This method of rating originality on a five-point rating scale has also been used in a Japanese creativity study (Yamaoka and Yukawa, 2017). After rating the answers’ originality, an average score was calculated as the score of each object. Then, the average score of the five objects, evaluated by three judges, was calculated as the task score. ICC for the originality score of all objects ranged from 0.56 to 0.81 for inter-rater reliability (Table 2). ICC values for each task type, averaging the value of the five objects, ranged from 0.65 to 0.72 and showed good reliability (Cicchetti, 2001). In the study by Hass et al. (2018), in order to promote reliability across the AUT prompts often used by researchers as measures of divergent thinking, generalizability and dependability studies were conducted for the average-rating system to facilitate feasibility and validity of ratings performed by laypeople. Results showed that good reliability can be achieved on the divergent thinking task (AUT) using the average-rating system and a specific number of items and raters. Figure 2 of their study illustrates that with three raters, four alternative uses objects would suffice to achieve sufficient generalizability if studies are conducted with the average-rating system, as we did in the present study. Each type of AUT used in the present study consisted of five objects and was judged by three raters.

**Table 2.**
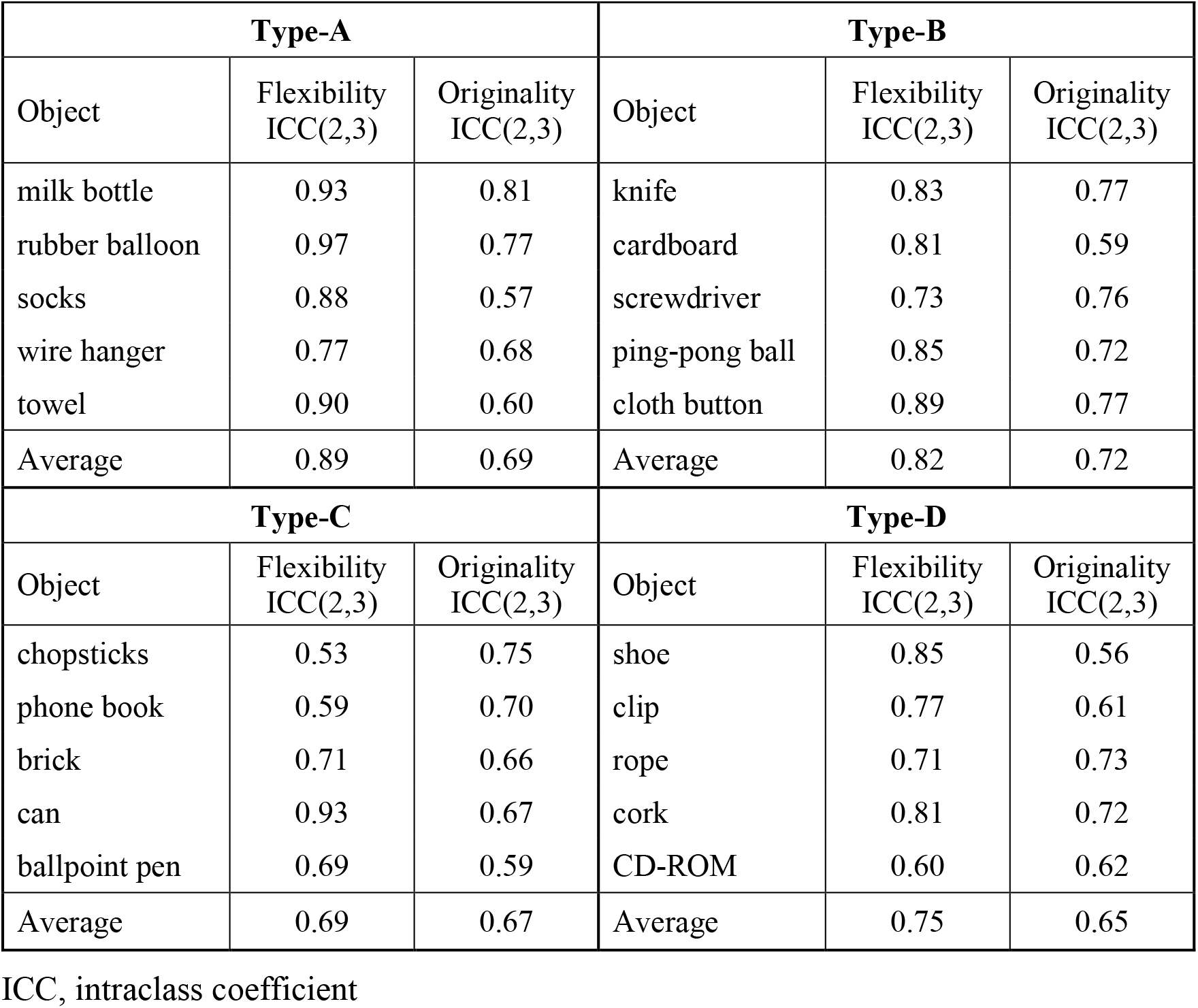
Intraclass coefficient (ICC) values for flexibility and originality scores of all objects

### 2.5 Experimental procedure

All participants underwent both Real-Stim and Sham-Stim on two separate days, each for 20 minutes, with an interval ≥three days between experimental sessions. The experiment consisted of a stimulation session and two EEG recording sessions before and after, with a 5–10-minute break between sessions (Fig. 1). The EEG/tDCS device attachment/removal and sleepiness questionnaires were conducted during the break. During the EEG recording session, we measured EEG for 15 minutes while the participant was answering the AUT, and for 1 minute during resting state, with eyes open, before the AUT. During the tDCS session, the participants watched a driving scene of a car taken by the drive recorder and verbally answered when they noticed a change in driving scene, in order to prevent sleep. The order of tDCS types (Real-Stim and Sham-Stim) and the order of AUT types (A, B, C and D) were counterbalanced among participants to take into account the order effects.

**Fig. 1.**
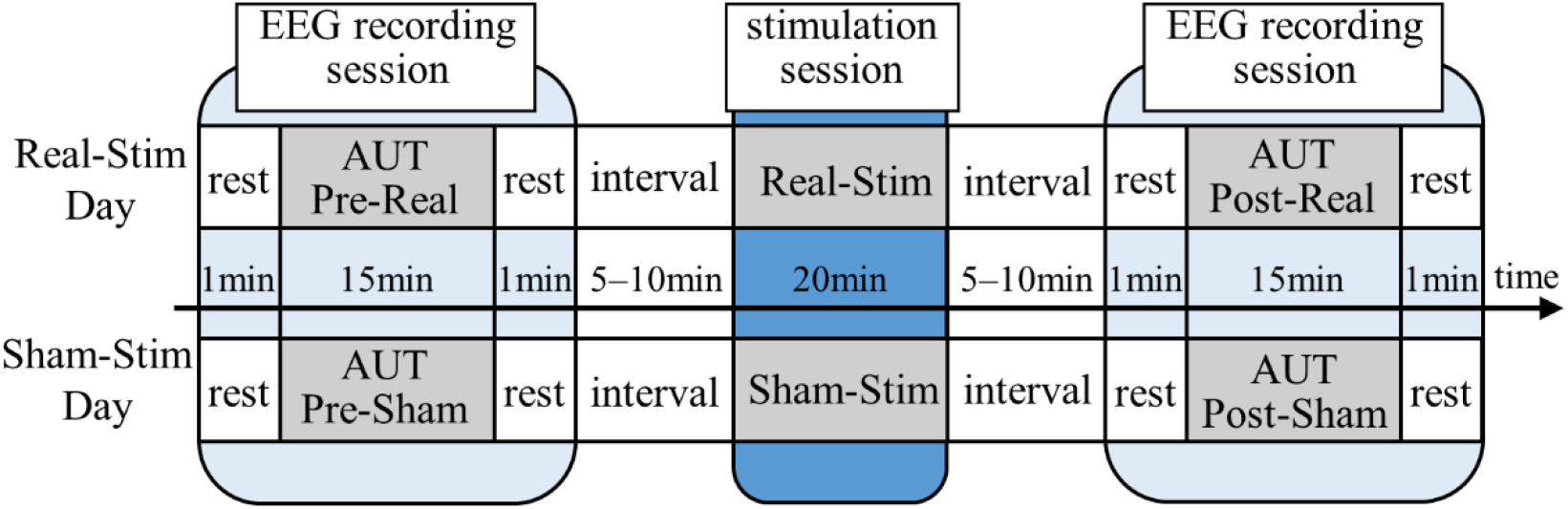
**Experimentalparadigm.** Real-Stim, real direct current stimulation; Sham-Stim, sham stimulation.

### 2.6 EEG measurement and preprocessing

EEG was recorded using the BrainAmp DC (Brain Products) and Brain Vision Recorder (Brain Products) from 32 places over the whole head according to the International 10-20 system (Fp1/2, F7/8, F3/4, Fz, FT9/10, FC 5/6, FC 1/2, T 7/8, C 3/4, Cz, CP 5/6, CP 1/2, TP 9/10, P7/8, P3/4, Pz, O1/2, and Oz) using the ActiCap (Brain Products) with silver-silver chloride active electrodes. During EEG measurements, the electrode located at Fpz was used as ground and the one located at FCz as the system reference. The sampling frequency was 500 Hz, the time constant was 10 s, and the high cut filter was 1000 Hz.

The data were analyzed offline using MATLAB 2016b and EEGLAB (Delorme and Makeig, 2004) version 14.1.b. Recorded EEG signals were bandpass-filtered between 0.5 Hz and 100 Hz using the FIR filter (EEGLAB function “pop_eegfiltnew.m”). Power line fluctuations at 50 Hz were removed using a notch filter (EEGLAB function “pop_cleanline.m”). Electrooculographic (EOG) and electromyographic (EMG) artifacts were removed using the Automatic Subspace Reconstruction (ASR) method (EEGLAB function “clean_rawdata.m” (Mullen et al., 2015; Chang et al., 2018)). The ASR threshold was set at 15 standard deviations based on the recommended value range from the EEGLAB website (Makoto’s preprocessing pipeline), with all other parameters turned off. EEG data epochs during the AUT (5 objects × 2 min) were extracted. The EEG data were re-referenced to a common average reference.

### 2.7 Connectivity analysis

Exact low resolution brain electromagnetic tomography (eLORETA, LORETA Key software version 20181107 (Pascual-Marqui et al., 1994)) was used to compute the connectivity strength between estimated cortical signals from multichannel head-surface EEG data (Pascual-Marqui et al., 2011). We chose eLORETA because of its accurate estimation of the intracortical distribution of current source density, effectively reducing the effects of volume conduction (Pascual-Marqui et al., 2014b). LORETA validation for localization agreement with multimodal imaging techniques has been reported in several studies (Worrell et al., 2000; Mulert et al., 2004; Zumsteg et al., 2005). Further, previous studies reported that eLORETA can be used to estimate deep brain source activities, including hippocampus and anterior cingulate cortex (Pizzagalli et al., 2004; Cannon et al., 2005; Lamm et al., 2014; Auerbach et al., 2015). The head model and electrode coordinates were based on the Montreal Neurologic Institute average MRI brain (MRI152) (Fonov et al., 2011). The solution space was restricted to the cortical gray matter (6239 voxels at 5×5×5 mm spatial resolution). In order to estimate connectivity between core regions of the large-scale brain networks (DN subsystems, Salience, and ECN), 11 cortical regions of interest (ROIs) were selected (Table 3).

**Table 3.** List of 11 cortical regions of interest (ROIs) selected for connectivity analyses

| ROI | MNI Coordinates |  |  | Anatomical Region | Brain Network |
| --- | --- | --- | --- | --- | --- |
|  | x | y | z |  |  |
| 1 | 0 | 55 | 10 | frontal lobe, medial prefrontal cortex (mPFC) | DN<sub>CORE</sub> |
| 2 | 0 | -50 | 25 | limbic lobe, posterior cingulate cortex (PCC) | DN<sub>CORE</sub> |
| 3 | -45 | -45 | 35 | left parietal lobe, inferior parietal lobule (l-IPL) | DN<sub>CORE</sub> |
| 4 | 45 | -50 | 35 | right parietal lobe, inferior parietal lobule (r-IPL) | DN<sub>CORE</sub> |
| 5 | -20 | -25 | -10 | left limbic lobe, hippocampus (l-HF) | DN<sub>MTL</sub> |
| 6 | 20 | -25 | -10 | right limbic lobe, hippocampus (r-HF) | DN<sub>MTL</sub> |
| 7 | -55 | -15 | -20 | left temporal lobe, lateral temporal cortex (l-LTC) | DN<sub>SUB3</sub> |
| 8 | 55 | -15 | -20 | right temporal lobe, lateral temporal cortex (r-LTC) | DN<sub>SUB3</sub> |
| 9 | 0 | 30 | 20 | limbic lobe, anterior cingulate cortex (ACC) | Salience |
| 10 | -40 | 40 | 25 | left frontal lobe, dorsolateral prefrontal cortex (l-DLPFC) | ECN |
| 11 | 40 | 40 | 25 | right frontal lobe, dorsolateral prefrontal cortex (r-DLPFC) | ECN |

DN, default mode network; DN<sub>CORE</sub>, core DN subsystem; DN<sub>MTL</sub>, medial temporal lobe-centered DN subsystem; DN<sub>SUB3</sub>, third DN subsystem; Salience, salience network; ECN, executive control network

Functional connectivity (FC) was analyzed using Lagged Phase Synchronization (LPS). LPS has been widely used to investigate electrophysiological connectivity (Canuet et al., 2011; Hata et al., 2016; Imperatori et al., 2017). Since detailed information on eLORETA LPS has been previously described (Pascual-Marqui et al., 2011), here we summarize the method. LPS performs a discrete Fourier transform of two signals followed by normalization to evaluate the similarity of signals in a specific frequency band. The equations representing LPS between signals *x* and *y* are:

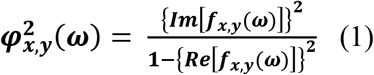

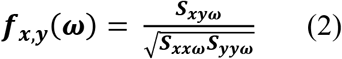

where *S*_*xxω*_, *S*_*xyω*_, and *S*_*yyω*_ represent complex valued covariance matrices, and *f*_*xy*_ is the complex-valued coherence. LPS is considered to accurately represent FC, as it excludes the instantaneous phase synchronization due to non-physiological artifacts and volume conduction. The FC between all pairs of ROIs was computed for five frequency bands: δ (0.5–3.5 Hz); θ (4–7.5 Hz); α (8–12.5 Hz); β (13–30 Hz); and γ (30.5–60 Hz).

Effective directional connectivity (EC) was assessed with isolated effective coherence (iCoh). This EC analysis is performed at the source level, so it requires EEG source localization (Grech et al., 2008; Jatoi et al., 2014). Since details of the iCoh method have been previously described (Pascual-Marqui et al., 2014b, 2014a), we briefly summarize it. The equation representing iCoh is:

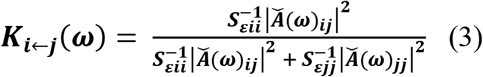

where *K*_*i*←*j*_(*ω*) is the iCoh value at a given frequency *ω* between ROI *i* and *j*, the arrow indicating that *j* influences *i*. 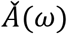 is the discrete Fourier transform matrix derived by least square fitting of the MVAR model of order *p* (estimated by the Akaike information criterion). *S*_*ɛ*_ is the covariance matrix of the residual errors of the MVAR model. The LORETA software is able to automatically compute all parameters in this equation, producing an iCoh spectrum as output, when provided with EEG data as input (Pascual-Marqui et al., 2014a). In our case, the optional parameter *p* of the MVAR model was set to 8. The EC between all pairs of ROIs was computed for all frequencies in the 0-60 Hz range. By averaging the 5 iCoh-value for the 5 epochs corresponding to the 5 everyday objects composing an AUT, one mean iCoh-value was calculated for each participant and each of the four conditions (Pre/Post Real-Stim and Pre/Post Sham-Stim).

## 3 Results

### 3.1 Creativity score change (Real-Stim vs Sham-Stim)

There was no significant difference between Real-Stim and Sham-Stim types for changes in all creativity scores: fluency: *t*(13) = −0.23, *dz* = 6.10e-2, *p* = 0.82 (M_Real_ = 0.54, SD_Real_ = 0.99; M_Sham_ = 0.63, SD_Sham_= 0.94); flexibility: *t*(13) = −0.33, *dz* = 8.85e-2, *p* = 0.75 (M_Real_ = 0.28, SD_Real_ = 1.04; M_Sham_ = 0.40, SD_Sham_ = 1.01); originality: *t*(13) = 0.34, *dz* = 9.03e-2, *p* = 0.74 (M_Real_ = −0.02, SD_Real_ = 0.22; M_Sham_ = −0.06, SD_Sham_ = 0.20).

### 3.2 FC changes during creative thinking (Post-Stim vs Pre-Stim)

On Real-Stim, an increased δ band FC between the posterior cingulate cortex (PCC) and r-IPL (*t*(13) = 4.50, *dz* =1.20, raw *p* = 5.92e-4, corrected *p* < 0.05, M_Post-Stim_ = 3.73e-2, SD_Post-Stim_= 5.15e-4; M_Pre-Stim_ = 1.59e-2, SD_Pre-Stim_= 9.88e-5) was observed (Fig. 2). No significant changes were observed in the other frequency bands. On Sham-Stim, there were no significant differences in FC in any frequency band. There was no significant difference in the paired stimulation type comparison.

**Fig. 2.**
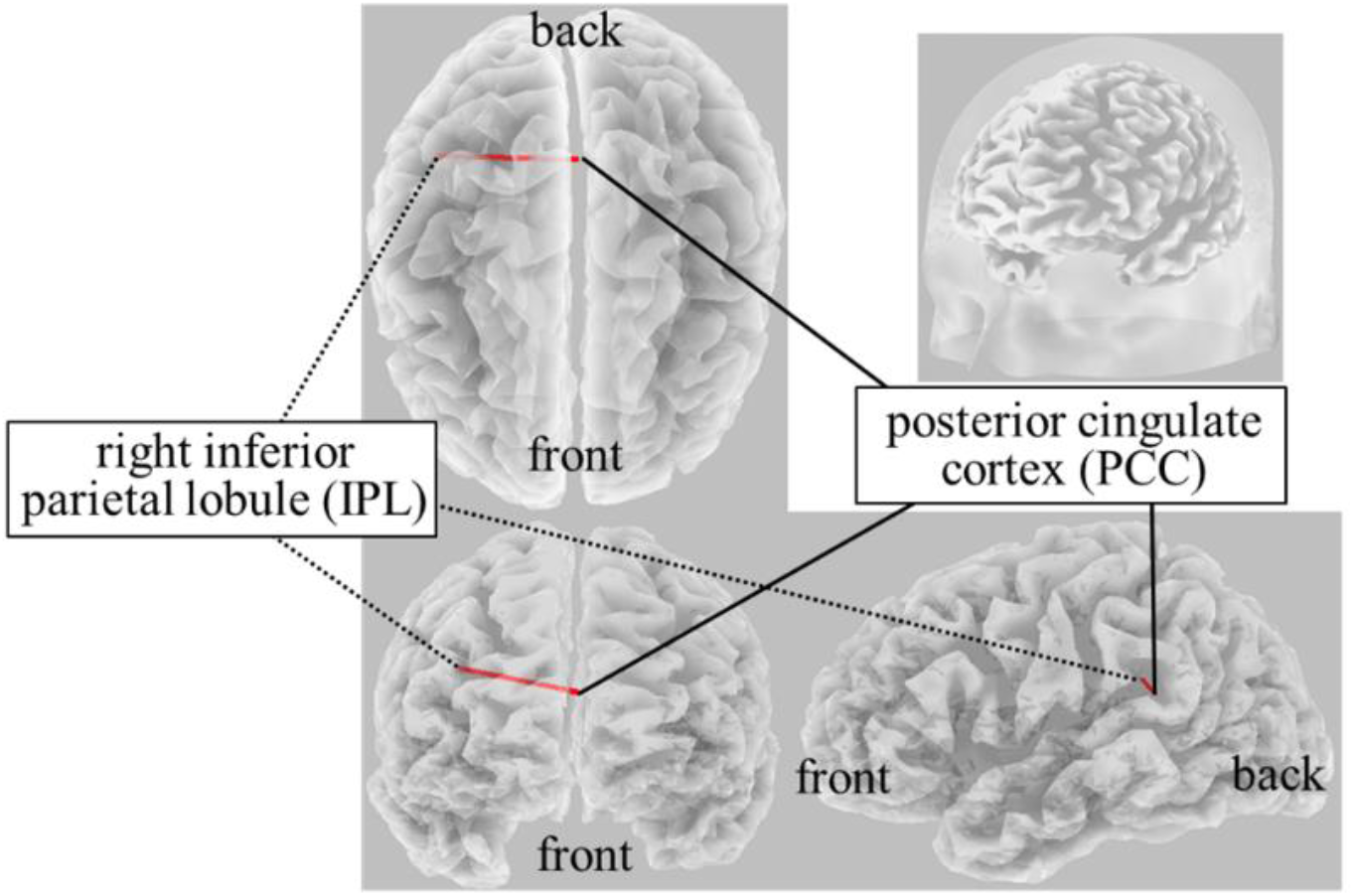
**Result of eLORETA for the comparison of functional connectivity in the delta frequency band between pre- and post-tDCS.** The red line indicates the connection showing increased LPS after tDCS (corrected p < 0.05).

### 3.3 Correlation between FC changes and creative thinking score change

On Real-Stim, the change in flexibility score was strongly and positively correlated with changes in the following: (i) the δ-band FC between the medial prefrontal cortex (mPFC) and the left lateral temporal cortex (l-LTC) (Spearman’s *rho* = 0.815, raw *p* = 6.24e-4) (Fig. 3); (ii) the α-band FC between the right lateral temporal cortex (r-LTC) and r-IPL (Spearman’s *rho* = 0.829, raw *p* = 3.97e-4) (Fig. 4). There was no correlation between these connectivities and the other creativity scores (fluency and originality). No significant correlations were found between changes in creativity scores and FC changes in the other frequency bands. On Sham-Stim, there was no significant correlation between FC changes and changes in creativity score.

**Fig. 3.**
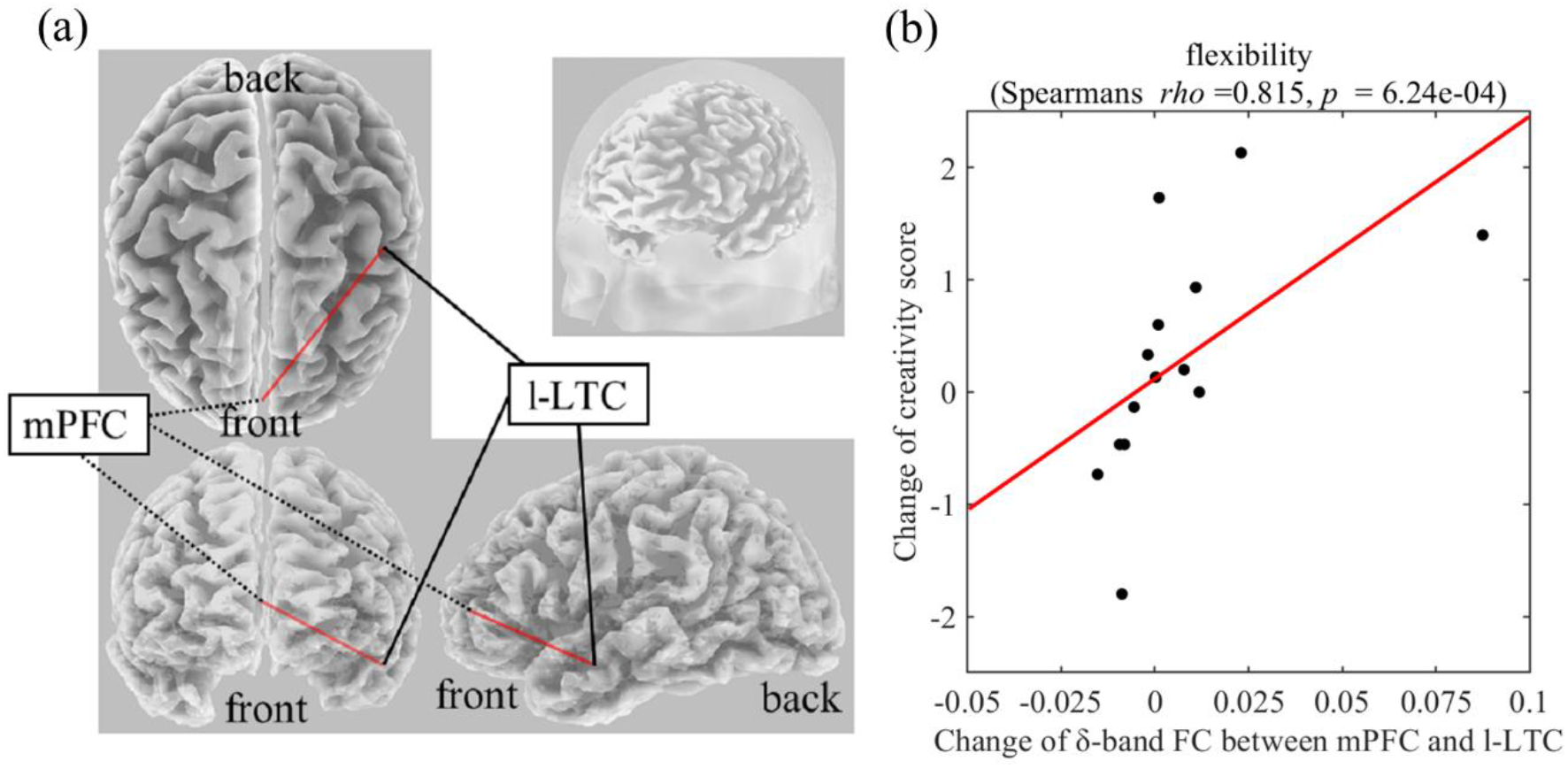
**(a) Functional connectivity between mPFC and l-LTC in the δ frequency band (b) Correlation between the change in δ-band functional connectivity between mPFC and l-LTC and the change in flexibility score upon Real-Stim.** The scatterplot shows the data of each participant. The red line indicates the least-squares regression line. FC, functional connectivity; mPFC, medial prefrontal cortex; l-LTC, left lateral temporal cortex.

**Fig. 4.**
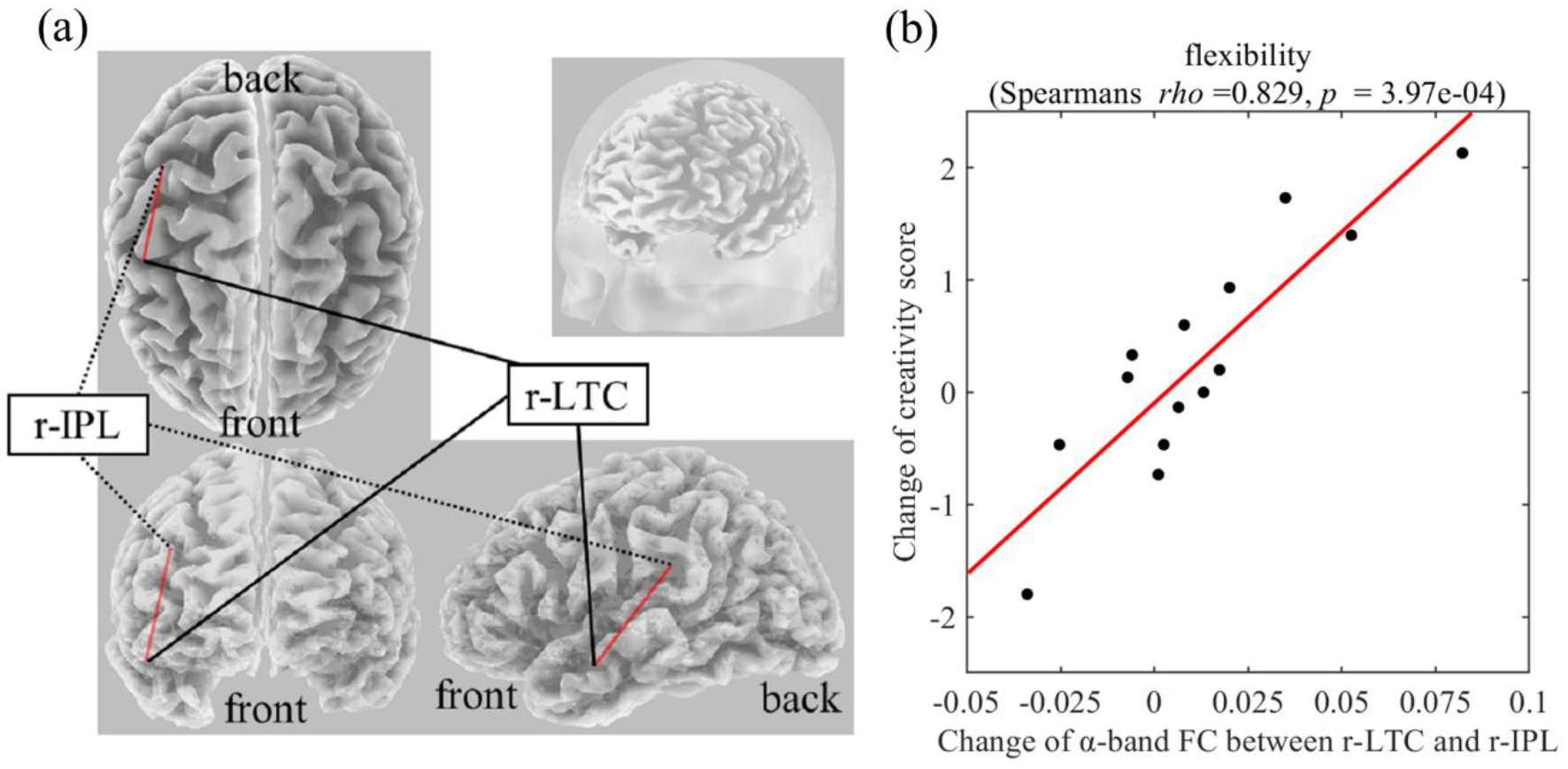
**(a) Functional connectivity between r-LTC and r-IPL in the α frequency band (b) Correlation between the change in α-band functional connectivity between r-LTC and r-IPL and the change in flexibility score upon Real-Stim.** The scatterplot shows the data of each participant. The red line indicates the least-squares regression line. FC, functional connectivity; r-LTC, right lateral temporal cortex; r-IPL, right inferior parietal lobule.

### 3.4 EC change during creative thinking (Post-Stim vs Pre-Stim)

There was no significant difference between Post-Stim and Pre-Stim in both stimulation types (Real-Stim and Sham-Stim). However, in the paired stimulation type comparison, significantly decreased low γ-band (29 Hz–37Hz, peak at 33 Hz) flow from l-LTC to r-IPL was observed (at the peak, 33 Hz; *t*(13) = −4.91, *dz* =1.31, raw *p* = 2.84e-4, corrected *p* < 0.05, M_Real-Stim(post-pre)_ = −7.62e-4, SD_Real-Stim(post-pre)_ = 7.58e-4; M_Sham-Stim(post-pre)_ = 4.20e-4, SD _Sham-Stim(post-pre)_= 9.28e-4) (Fig. 5). No significant changes were observed in the other flows between ROIs.

**Fig. 5.**
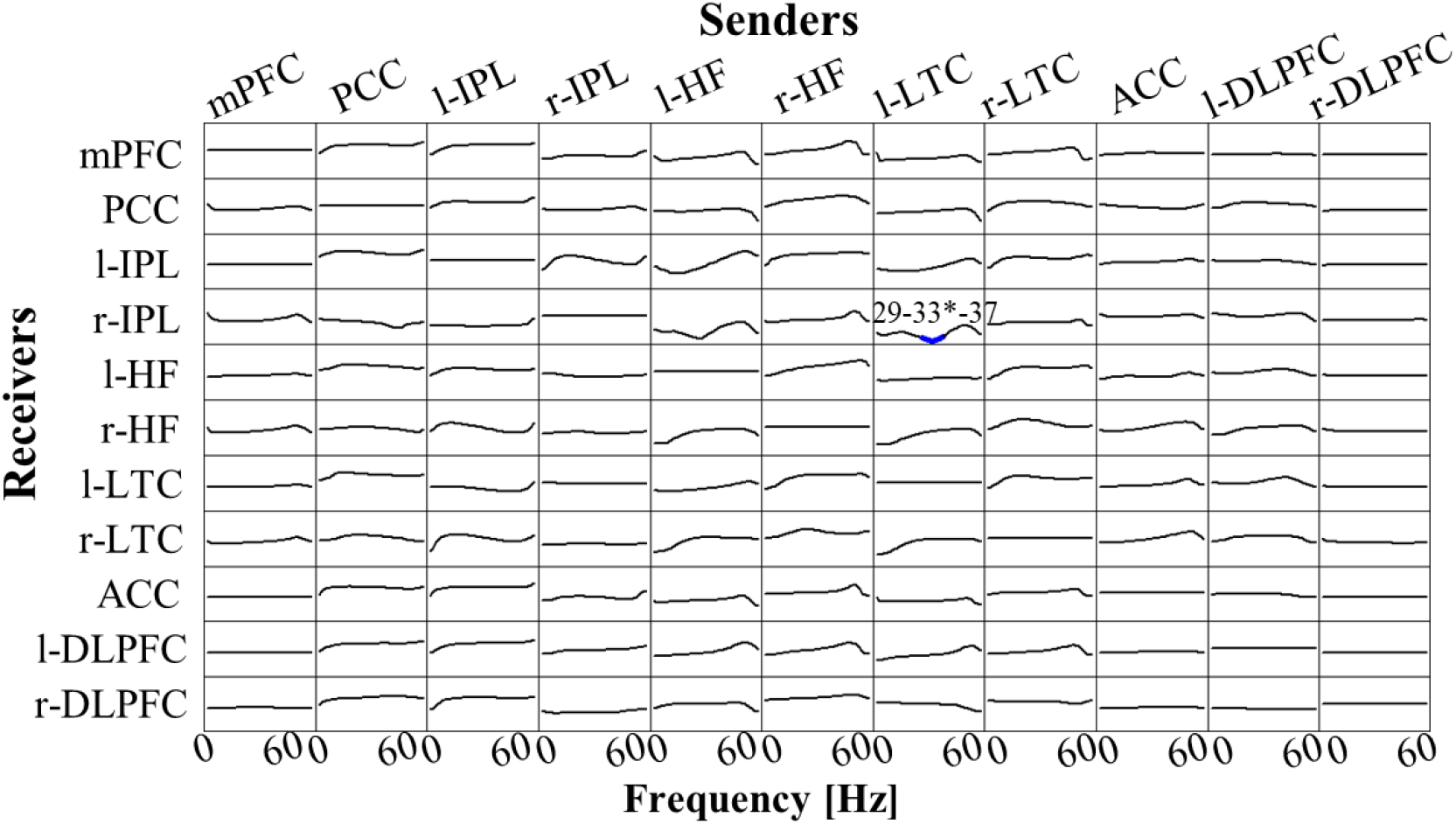
**t-statistic for the comparison of Real-Stim (post-pre) and Sham-Stim (post-pre) isolated effective coherence (iCoh) during creative thinking for 14 participants, in 11 regions of interest (ROIs):** mPFC, medial prefrontal cortex; PCC, posterior cingulate cortex; l-IPL/r-IPL, left and right inferior parietal lobule; l-HF/r-HF, left and right hippocampus, l-LTC/r-LTC, left and right lateral temporal cortex; ACC, anterior cingulate cortex; l-DLPFC/r-DLPFC, left and right dorsolateral prefrontal cortex. Frequency axis: 0-60 Hz. Corrected p = 0.05 corresponds to a t-threshold of 4.54, with the vertical axis spanning from −5.0 to +5.0. Blue (red) color indicates significantly larger values in Sham-Stim (Real-Stim). The most significant oscillation is indicated with a superscript “*”.

## 4 Discussion

Here, we investigated the effects of tDCS (anode over l-DLPFC and cathode over r-IPL) on brain networks during creative thinking and the causal relationships between the changes in connectivity of the large-scale brain network related to creative thinking, and changes in creative performance. We found that applying tDCS increased δ-band FC between r-IPL (DN_CORE_) and PCC (DN_CORE_) and decreased the low γ-band EC from l-LTC (DN_SUB3_) to r-IPL (DN_CORE_) during divergent thinking. Creativity performance (fluency, flexibility and originality) was not significantly affected by the stimulation, but (i) the change of δ-band FC between mPFC and l-LTC; and (ii) the change of α-band FC between r-IPL (DN_CORE_) and r-LTC (DN_SUB3_) induced by tDCS are reflected in a change in creative thinking flexibility. The present study is the first to investigate the effects of tDCS on brain networks (EEG-index) related to creative thinking, and their causal relationships with creative performance.

It has been suggested that anodic/cathodic stimulation for l-DLPFC and r-IPL affects DN activity at the resting state and mind-wandering (Axelrod et al., 2015; Kajimura and Nomura, 2015; Kajimura et al., 2016). The present study extended the suggestion to divergent thinking. The right IPL is known to causally affect other DN regions (Di and Biswal, 2014). Applying cathodal stimulation could have decreased the neuron excitability under the right IPL and causally modulated the other regions of DN. The l-DLPFC, the core region of the ECN, is known to be flexibly coupled with DN during divergent thinking and be involved in attentional shift (Wager et al., 2004), flexible switching between semantic categories (Kleibeuker et al., 2013), and maintenance of internally generated thoughts (Burgess et al., 2003). Therefore, the anodal stimulation on the l-DLPFC (ECN) may have affected the DN during AUT and supported the attention shift from the original use of objects to self-generated thought and flexible switching between semantic categories.

The r-IPL and PCC both belong to the DN_CORE_, which acts as a hub within the DN and contributes to internal oriented cognition (Christoff et al., 2016). The r-IPL and PCC are both activated when we remember past events or imagine future ones (Addis et al., 2007; Abraham et al., 2008). A previous fMRI study also reported that the FC between right PCC and r-IPL significantly increased during AUT compared with a control task consisting in generating typical properties of everyday objects (Beaty et al., 2015). Therefore, it is possible that during AUT participants remember past experiences or imagine future events in which the everyday object is used for alternative purposes. In the present study, effects were not manifested in the creativity score changes but the present findings indicate that the increase in FC between r-IPL and PCC induced by tDCS may causally facilitate the initial creative thinking process, namely the spontaneous generation of novel ideas, during AUT. The low γ-band flow from l-LTC (DN_SUB3_) to the r-IPL (DN_CORE_) significantly decreased in Real-Stim compared to Sham-Stim. This effective connectivity has not been reported in the previous creativity research, but considering that the change in FC between (i) mPFC(DN_CORE_) and l-LTC(DN_SUB3_); (ii) r-IPL (DN_CORE_) and r-LTC (DN_SUB3_) showed a strong positive correlation with the change in flexibility score, this decreased flow may have causally affected these FCs within the DN and then affected the flexibility of the output.

Correlation results of the present study indicate that the change of δ-band FC between mPFC and l-LTC positively affects the flexibility of divergent thinking. The LTC is a core brain region of DN_DUB3_ and the exact anatomical region selected as ROI in this study is the middle temporal gyrus (MTG). The left-MTG plays a key role in semantic processing (Cappa, 2008; Whitney et al., 2011; Abraham, 2014), and has been suggested to be associated both with the acquisition of semantic knowledge and the retrieval of different types of semantic information such as semantic functional knowledge (abstract properties, as function and context of use) and semantic action-related knowledge (motor-based knowledge of object utilization) (Vandenberghe et al., 1996; Phillips et al., 2002; Maguire and Frith, 2004; Canessa et al., 2008). Furthermore, previous research has shown that generating action words to visually presented objects activates the left-MTG (Martin and Chao, 2001). On the other hand, it has been suggested that the mPFC is involved in retrieval of long-term memories and future thinking (Takashima et al., 2006; Gais et al., 2007; Quinn et al., 2008; Euston et al., 2012). A previous study reported that the anterior mPFC was more activated during personal future, personal past and non-personal future thinking than during non-personal past thinking (Abraham et al., 2008). Furthermore, Spalding et al. (2018) suggested that the ventral mPFC plays key roles in the application of existing knowledge to novel circumstances. Together, the FC between mPFC and l-LTC during divergent thinking (e.g., AUT) might play a key role in generating action words of alternative use by “flexibly” recombining semantic knowledge of the everyday object referencing past personal memory. FC with mPFC as a node has been proposed in association to the individual difference in creativity measured by divergent thinking tasks (Takeuchi et al., 2012; Wei et al., 2014). A previous functional imaging study reported that creativity measured by divergent thinking task was significantly positive correlated with resting-state FC between mPFC and left MTG (Wei et al., 2014). These previous findings indicate the involvement of FC between mPFC and l-LTC in creativity while the present finding regarding a causal relationship extends the possibility that this FC causally facilitates the flexibility of divergent thinking.

In addition, the change of α-band FC between r-IPL (DN_CORE_) and r-LTC (DN_SUB3_) also positively affected the flexibility of divergent thinking. Several previous results support the important role of the right MTG for insight (Subramaniam et al., 2009; Cranford and Moss, 2011; Sakaki and Niki, 2011; Wu et al., 2013). When people encounter words, they think about related information (Jung-Beeman, 2005). “Semantic activation” provides access to semantic representations, activation features, and first order associations of input words. This semantic activation depends on Wernicke’s areas in both hemispheres, and especially the posterior middle and superior temporal gyrus. The left hemisphere strongly activates small and focused semantic fields containing information closely related to the dominant meaning of the input words. In contrast, the right hemisphere weakly activates large diffuse semantic fields, containing distant and unusual semantic features unrelated to input words, providing coarse interpretation (Jung-Beeman, 2005). During AUT in this study, participants encountered a word representing an everyday object and were asked to think about as many alternative uses as possible. It is possible that participants were searching for semantically distant related words. Considering the role of r-LTC (DN_SUB3_) in distant semantic activation and the role of r-IPL (DN_CORE_) in internal oriented cognition, the strength of the functional connection between these two DN regions may causally affect the flexibility of creative thinking.

This study has some limitations. First, the sample was small and biased, for instance, in terms of sex and age range, which may affect the robustness of results. Second, there is no control stimulation in terms of polarity or region. Therefore, it is difficult to definitively link our findings to the specific target regions stimulated. Third, the size of the electrodes used for tDCS was relatively large, making it difficult to perform exact and focal stimulation of the l-DLPFC and r-IPL. Thus, the current results should be replicated using more targeted neurostimulation techniques, such as High-Definition tDCS (Edwards et al., 2013). Fourth, to avoid sleepiness and influencing the results of the verbal creativity task, participants watched a video content least relevant to the everyday object used in the AUT during the tDCS session. However, there is certainly no denying the possibility that this procedure led to the activation of visuo-spatial circuits and evoked visual imagery during Post-Stim AUT. In our study, it is difficult to determine how much of an effect of activating these visuo-spatial circuits on the task performance and on other brain networks related to creative thinking, but we can speculate that the activation effect was manifested in the results of both the real-stim and sham-stim conditions. Fifth, we did not have tDCS sessions within the same time span for every participant (three or more days between the experimental sessions). However, previous studies have shown that increased cortical excitability lasts for 90 minutes after a 13-minute anodal stimulation and reduced cortical excitability lasts for 60 minutes after a 9-minute cathodal stimulation. Therefore, ≥3 days between Real-Stim and Sham-Stim was likely sufficient to prevent the effects of the first tDCS session from influencing the second. Sixth, we selected and analyzed only certain brain regions, but others may be involved in creative thinking. However, we identified some mechanism underlying creative thinking after tDCS.

## 5 Conclusions

The current study used connectivity analyses to investigate the effects of tDCS on brain networks during divergent thinking and the associated change in creativity performance. Our findings provide new evidence regarding the neural mechanism of creative thinking, particularly flexibility. Future research is needed to clarify the causal relationship between mechanism and creative performance, with the aim of devising methods for the enhancement of creativity.

## 7 Author contributions

KK, KU, and ZL contributed conception and design of the study. MN supervised the study. KK and ZL conducted the experiment and analyzed the data. KK wrote the first draft of the manuscript. All authors contributed to manuscript revision, read and approved the submitted version.

## 8 Acknowledgments

This research did not receive any specific grant from funding agencies in the public, commercial, or not-for-profit sectors. We would like to thank Editage (www.editage.com) for English language editing.

We presented the preliminary results of the present study in the following conferences: The 9th International IEEE EMBS Conference on Neural Engineering and LIFE 2019 (Internal conference). Although we analyzed the same experimental data for these conferences, after receiving feedback from other researchers, we changed the analysis method and added new contents in this article.

